# Protein Arginine Deiminase 4 Antagonizes Methylglyoxal-induced Histone Glycation

**DOI:** 10.1101/826818

**Authors:** Qingfei Zheng, Adewola Osunsade, Yael David

**Affiliations:** Chemical Biology Program, Memorial Sloan Kettering Cancer Center, New York, New York 10065, United States; Tri-institutional PhD Program in Chemical Biology, New York, New York 10065, United States; Department of Pharmacology, Weill Cornell Medicine, New York, New York 10065, United States; Department of Physiology, Biophysics and Systems Biology, Weill Cornell Medicine, New York, New York 10065, United States

**Keywords:** protein arginine deiminase 4, histone modification, methylglyoxal-glycation, metabolism, epigenetics

## Abstract

Protein arginine deiminase 4 (PAD4) facilitates the post-translational citrullination of the core histones H3 and H4. While the precise epigenetic function of this modification has not been resolved, it was shown to associate with general chromatin decompaction and to compete with arginine methylation. Recently, we showed that histones are subjected to methylglyoxal (MGO)-induced glycation on nucleophilic side chains, particularly arginines, under metabolic stress conditions. These non-enzymatic adducts change chromatin architecture and the epigenetic landscape by competing with enzymatic modifications. Here we report that PAD4 antagonizes histone MGO-glycation by protecting the reactive sites with oxygen substitution, as well as by converting already-glycated arginine residues into citrulline. Moreover, we show that similar to the deglycase DJ-1, PAD4 is overexpressed and histone citrullination is upregulated in breast cancer tumors, suggesting an additional mechanistic link to PAD4’s oncogenic properties.

**Significance:** Metabolic syndromes and diabetes increase the risk for certain diseases such as cancer. However, the mechanism behind this correlation is poorly understood. Methylglyoxal (MGO), a reactive dicarbonyl sugar metabolite found in cells under metabolic stress, can non-enzymatically modify arginine and lysine residues in histone proteins, making it a new epigenetic marker linking metabolism and disease. Histone MGO-glycation induces changes in chromatin architecture and the epigenetic landscape, and abrogates gene transcription. In this study, we found that protein arginine deiminase 4 (PAD4) exhibits dual functions to antagonize histone MGO-glycation: removing glycation adducts from arginines and converting the unmodified side chains into citrulline, which protects them from undergoing glycation. This unprecedented biochemical mechanism demonstrates a potential function of PAD4 in cancer cells.

---

In eukaryotes, nucleosomes are the fundamental unit of chromatin, composed of DNA and histone proteins (1). Post-translational modifications (PTMs) on histones, including acetylation and methylation, are important in regulating chromatin structure and function during replication, transcription and DNA-damage (2, 3). It is speculated that the combinations of specific PTMs form a so called “histone code” (4) that is established through a network of crosstalk between enzymatic modifications and determines the transcriptional state of a specific genomic locus (5, 6). Citrullination, which occurs on arginine residues and involves the deimination of the guanidino group, reduces the net charge of the side chain and was generally shown to promote chromatin fiber decompaction (7), although specific sites have also been associated with gene repression (8). It was previously demonstrated that histone H3 arginine citrullination antagonizes its methylation by both blocking the modification sites and preventing the recruitment of the methyltransferases (9, 10). Histone arginine citrullination is executed by the calcium-dependent enzyme, protein arginine deiminase 4 (PAD4). PAD4 substrates include certain sites on both core and linker histones (11), and it was suggested to play a role in determining cellular pluripotency as well as in the DNA damage response (12). Although *PAD4* (*PADI4*) is a documented oncogene (13-16), the regulatory function of histone citrullination in pathological and physiological processes is still poorly understood (12).

Whereas the most well-characterized histone modifications are enzymatic, in the past few years it has become clear that histones are also prime substrates of non-enzymatic covalent modifications that induce changes in chromatin structure and function (17). We recently found that core histones are subjected to MGO-glycation and that these adducts change chromatin architecture, the epigenetic landscape, and transcription, and particularly accumulate in disease states (18). We determined that short or low-concentration exposure to MGO induces chromatin decompaction by compromising the electrostatic interactions of the histone tails with DNA, in a mechanism similar to acetylation. These adducts can rearrange and undergo crosslinking, which ultimately leads to chromatin fiber hyper-compaction. While MGO reacts with histones non-enzymatically, we found that DJ-1/PARK7 is a potent histone deglycase, preventing the accumulation of histone glycation *in vitro* and in cells (18). Since MGO rapidly reacts with the guanidino group of arginines, we tested the effect of MGO-glycation on cellular H3R8 methylation mark and found that MGO induces a reduction in H3R8me2 levels. As multiple arginine residues on histones (19), including H3R8, were shown to be substrates of PAD4, we hypothesized that citrullination, which alters the charges of amino acid side chains and reduces their reactivity against electrophiles, blocks MGO-glycation on arginines and vice versa. In this study, we addressed this hypothesis and unexpectedly found that beyond the direct competition between citrullination and MGO-glycation, PAD4 itself acts as a deglycase, mediating the conversion of arginine glycation adducts into citrulline (Figure 1).

**Figure 1.**
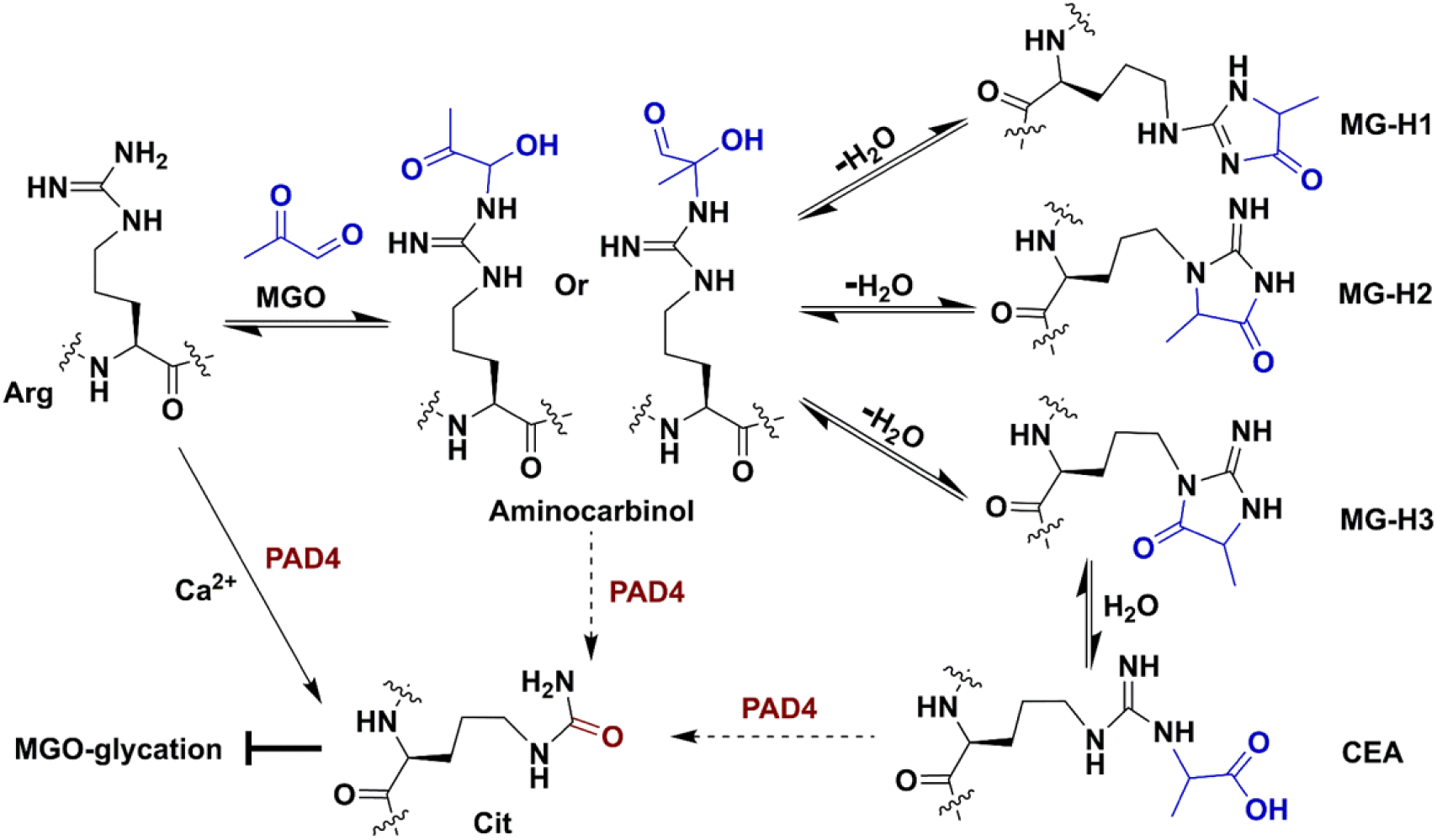
Overall schematic showing MGO-induced histone glycation and citrullination/deglycation activity of PAD4.

## Results

### PAD4-mediated Citrullination prevent histone from undergoing MGO-glycation

First, we aimed to analyze the biochemical mechanism of the crosstalk between histone citrullination and glycation. To do so, we purified recombinant PAD4 (Figure S1) and tested its activity *in vitro* on a range of substrates with increasing complexities, including free histone H3, nucleosome core particles (NCPs) and homododecameric (12-mer) nucleosomal arrays, which mimic the minimal chromatin fold (20). Our results indicate that PAD4 has increased reactivity towards more physiological substrates, with highest detected activity on 12-mer arrays (Figure S2), in a Ca^2+^-dependent manner (Figure S3). Next, we utilized unmodified and PAD4-citrullinated NCPs as substrates to test the direct competition between histone citrullination and MGO-glycation. Our results indicate that citrullination protects NCPs from undergoing MGO-glycation, since after a 12-hour MGO treatment the citrullinated NCPs were significantly less glycated compared to unmodified NCPs (Figure 2A). Moreover, the protective effect of PAD4-mediated citrullination against MGO-glycation is dose-dependent, (Figure S4).

**Figure 2.**
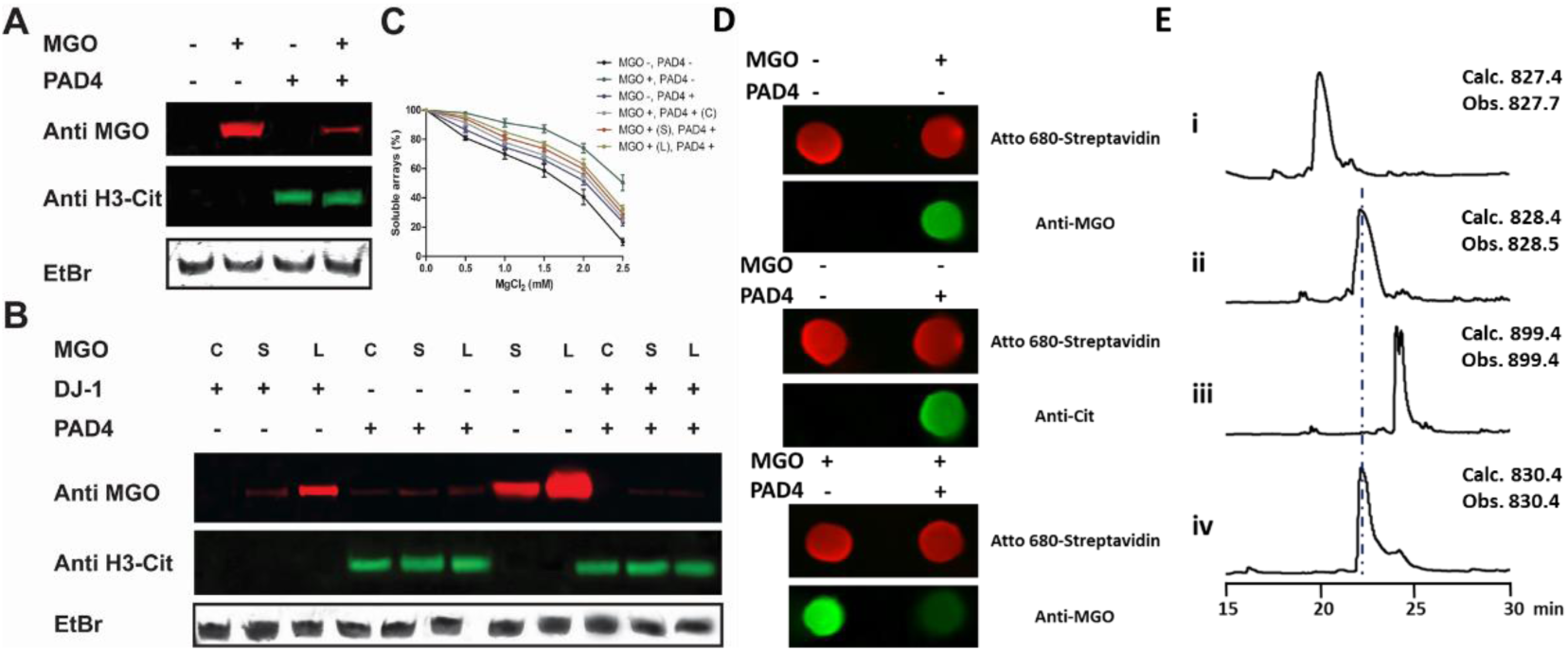
*In vitro* citrullination and glycation assays. (A) NCPs were first citrullinated by PAD4 followed by a 5 mM MGO treatment at 37 °C overnight. The reactions were separated on a native gel followed by a western blot analysis. (B) NCPs were pre-treated with 5 mM MGO for short (S, 6 h) or long (L, overnight) periods and then incubated with PAD4, or concurrently (C) incubated with both the enzyme and MGO for the same time. (C) PAD4 was incubated with glycated nucleosomal arrays after short (S) or long (L) treatment with MGO, or co-incubation (C). (D) Dot blot analysis of MGO-glycation and PAD4-mediated deglycation on H3 N-terminal peptide substrate. The H3 peptide sequence is NH_2_-ARTKQTARKSTGGKAPRK(Bio)A-COONH_2_, and the biotin signal (imaged by Atto 680-Streptavidin) was used as loading control. (E) LC-MS analysis of modified and unmodified H4 peptides (λ=214 nm): i) H4-R3(1-6) peptide, ii) H4-Cit3(1-6) peptide, iii) MGO-treated H4-R3(1-6), iv) MGO-glycated H4-R3(1-6) followed by PAD4 treatment in H2^18^O. The H4 synthesized peptide sequences are AcNH-SG<u>R</u>GK(Bio)G-COONH2 and AcNH-SG<u>Cit</u>GK(Bio)G-COONH_2_.

### PAD4 converts MGO-glycated histone arginines to citrullines

Next, we tested the reciprocal competition – that is, whether MGO-glycation protects NCPs from undergoing PAD4-mediated citrullination. To do so, we pre-treated NCPs with MGO and, after removing the excess MGO, used the glycated-NCPs as PAD4 substrates. The results unexpectedly indicated that PAD4 is still able to modify the glycated nucleosomes and that the citrullination is added at the expense of glycation, as evident by the decreased MGO-glycation and increased citrullination signals (Figure 2B). To test this newly identified histone deglycation activity of PAD4 and compare it to the one we recently identified in DJ-1 (18), we either added the enzymes concurrently with MGO to NCPs (C), or after a short (S) or a long (L) treatment with MGO (after removing unreacted MGO). The results indicate that DJ-1 is capable of removing MGO glycation from NCPs when added concurrently or after a short exposure to MGO. However, glycation was persistent following a long exposure, which allows the rearrangement of the glycation adducts into late-stage products (Figure 1 and 2B). In contrast to DJ-1 and the no-enzyme control, PAD4 was able to remove MGO-adducts and install citrulline, regardless of the incubation time (Figure 2B), suggesting it is active on both early- (aminocarbinol) and late-stage (carboxyethyl arginine, CEA) MGO adducts. Indeed, applying both DJ-1 and PAD4 resulted in the complete abolishment of the glycation (Figure 2B). To further validate the direct deglycase activity of PAD4, we performed a deglycation assay on biotinylated H3 and H4 N-terminal peptide substrates. The peptide was incubated with MGO, immobilized, washed and finally treated with PAD4. The reactions were analyzed by both dot blot (Figure 2D) and LC-MS (Figures 2E, S5 and S6), confirming that PAD4 directly removes MGO glycation adducts from the H3 and H4 tails *in vitro*. Finally, to examine the effect of PAD4 activity on chromatin compaction, we performed a Mg^2+^ precipitation analysis on 12-mer arrays substrate. This assay relies on magnesium inducing the aggregation and precipitation of the 12-mer fibers, so the less compacted the arrays are, the more Mg^2+^ is required to precipitate them. Our results demonstrate that histone citrullination decompacts the arrays to a lesser degree than MGO treatment, and that PAD4 rescues the majority of MGO-dependent chromatin decompaction for all incubation conditions (Figure 2C). This is in contrast to the deglycase DJ-1 (21-23), which rescues glycation-induced decompaction only under short or co-incubation conditions (Figure S7).

### PAD4 antagonizes histone MGO-glycation and regulates chromatin compaction

To investigate PAD4-mediated crosstalk between glycation and citrullination in native chromatin, we examined the impact of its expression on histone arginine citrullination, glycation and methylation in cells. As expected, PAD4 overexpression in 293T cells induces an increase in histone citrullination at the expense of arginine methylation (Figure 3A and S8). In analogy to our *in vitro* results, treating these cells with increasing amounts of MGO caused the accumulation of H3 and H4 glycation (in addition to crosslinking), which was suppressed by PAD4 overexpression (Figure 3A). To dissect the deglycation function of PAD4, we performed a pulse-chase experiment where cells were first treated with a gradient of MGO concentrations and then washed with fresh media. After a 6 hour-recovery, the PAD4 plasmid was transfected and cells were grown for additional 12 hours before final harvesting. Analysis of the histone samples from this experiment revealed that overexpression of PAD4 resulted in a decreased MGO glycation signal, suggesting PAD4 is actively removing MGO adducts from histones in cells (Figure S9). Moreover, a global chromatin compaction analysis by micrococcal nuclease (MNase) digestion revealed that PAD4 rescues MGO-induced chromatin decompaction (Figure S10). Together, these results establish that PAD4 directly affects chromatin compaction by regulating the epigenetic crosstalk between histone MGO-glycation, citrullination and methylation.

**Figure 3.**
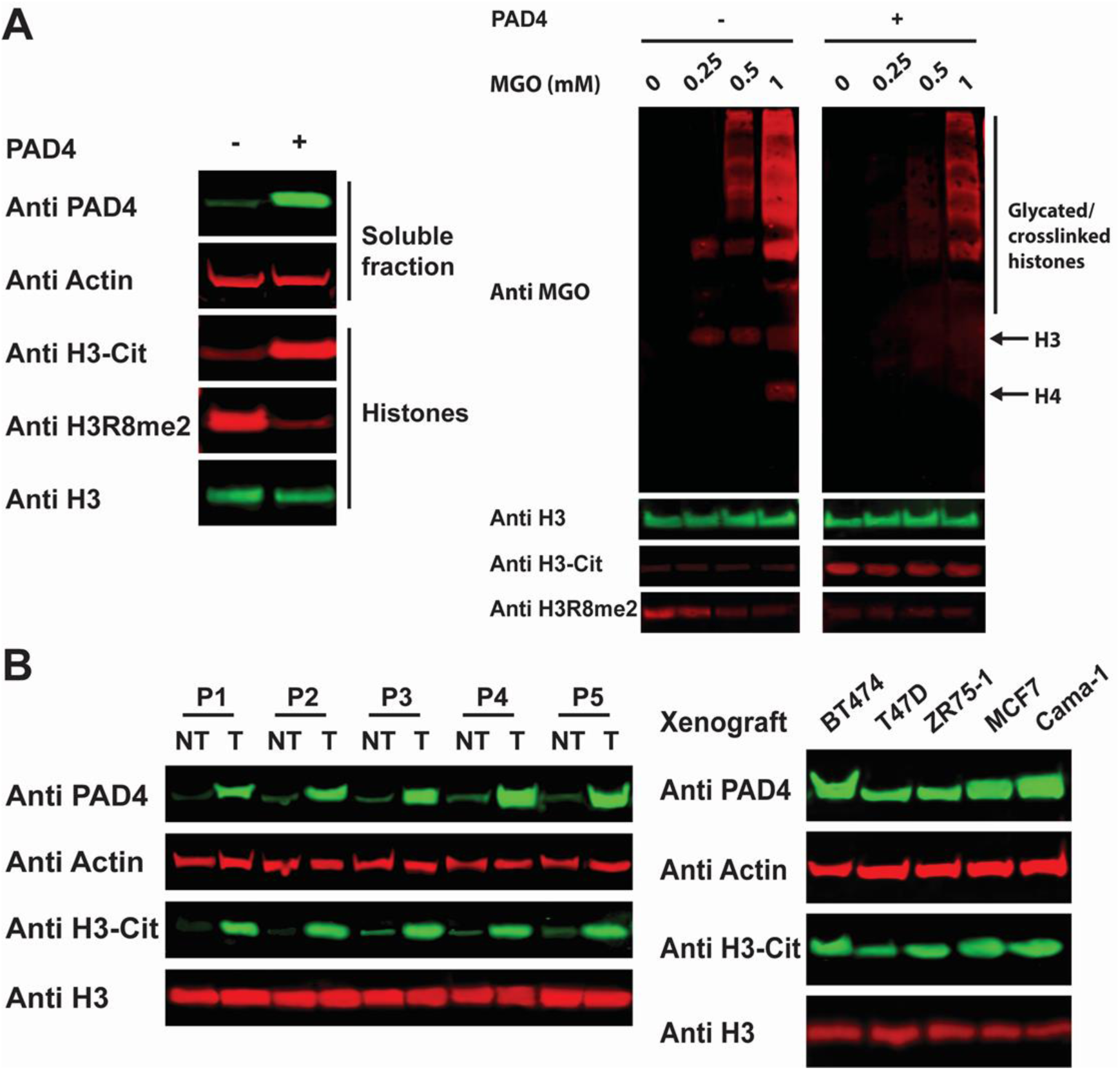
*In vivo* citrullination and glycation analyses. (A) Left: western blot analysis of both soluble and histone fractions of wild type and PAD4-overexpressing 293T cells. Right: western blot analysis of the histone fraction taken from the same cells that were treated with increasing concentration of MGO. (B) Western blot analysis of clinical tumor (T) and non-tumor (NT) samples from five different breast cancer patients (P1-P5) (left) or breast cancer xenografts (right).

### PAD4 is overexpressed in breast cancer

Previous studies demonstrated the overexpression of PAD4 in a variety of cancers, suggesting it could drive tumor pathogenesis through multiple potential mechanisms (12-16). Since we recently reported high levels of both histone MGO-glycation and DJ-1 expression in breast cancer (18), we utilized the same samples to analyze PAD4 and histone citrullination levels. As presented in Figure 3B, all patient tumor samples display substantial overexpression of PAD4 and increased levels of histone citrullination. In addition, xenograft tumor samples show diverse levels of PAD4 expression that correlate with degree of histone citrullination. In support, 293T cells treated with increasing amounts of MGO, which mimics the metabolic stress that exists in cancer cells, show better survival when overexpressing PAD4 but not the catalytically dead mutant C645S (Figures S11 and 4A) (9, 10). To investigate the impact of PAD4 on histone PTM crosstalk in breast cancer cells, MCF7 cells were pre-treated with the PAD4 inhibitor GSK484 (24) followed by treatment with increasing concentrations of MGO. The analysis, presented in Figure S12, shows that inhibition of PAD4 increases histone glycation and arginine methylation. Together, these data suggest a role for PAD4 in cancer proliferation through the regulation of chromatin structure and function.

## Discussion

Altogether, this study revealed a new crosstalk between histone glycation and citrullination mediated by PAD4. Our data support a model whereby PAD4 is capable of converting MGO-adduct intermediates into citrulline (Figures 1, 2 and S13), however, this new activity can proceed via different mechanisms (Figures S14 and S15). There are several implications for this newly identified function of PAD4 on MGO-glycated histones. First, it provides an additional pathway for the repair of pathophysiological histone glycation. We have previously shown that histone glycation is a modification that accumulates under metabolic stress, such as in highly-proliferating breast cancer tumors, and that it changes patterns of transcription and chromatin architecture. While we identified DJ-1 as a potent eraser of glycation, it is not surprising that cells developed multiple strategies to repair this severe chromatin damage. Indeed, we have shown that breast cancer patient samples contain massive overexpression of both DJ-1 (18) and PAD4 (Figure 3). Counterintuitively, these samples also have high levels of glycation, suggesting that the overexpression of DJ-1 and PAD4 is not sufficient to remove all the glycation adducts. The results we present here suggest that this could be due to the fact that DJ-1 can only erase early stage glycation products (18, 21-23), while PAD4 can specifically remove glycation modifications from arginine side-chains in a subset of proteins, including in the N-terminal residues of histones (12), which are key epigenetic regulators, by converting them to citrullines. Furthermore, other PAD enzymes, as well as PAD4 itself, may target glycation adducts in other substrates although this remains to be determined (12). We thus cannot rule out that additional repair mechanisms may exist to protect from or erase other glycation damage on histones or other cellular proteins.

In addition, our analysis reveals that breast cancer tumors contain high levels of histone citrullination, potentially at the expense of arginine glycation. It is possible that this is a general mechanism that exists in order to protect histones from damage and allow cancer to thrive despite metabolic stress. In that regard, both DJ-1 and PAD4 are proposed oncoproteins and targets of cancer therapy. There are several efficient DJ-1 and PAD4 inhibitors reported, some of which are in pre-clinical trials (25-27), raising the potential of combinational therapy targeting both enzymes simultaneously.

Both histone glycation at its early stages (18) and citrullination (7) induce chromatin decompaction, however glycation-induced decompaction seems to be more substantial and is suppressed by PAD4 (Figures 2D, S8 and S11). Unlike DJ-1, PAD4 is only active on arginine residues, which react preferentially with MGO relative to lysines (19). However, PAD4 is superior due to its dual function as a deglycase and protector, that is, it is able to both remove the modification and protect the residues from further damage. Although it was reported that free citrullines can be converted to arginines by argininosuccinate synthetase (ASS) and argininosuccinate lyase (ASL) in the citrulline-NO cycle (28), to date there is no report of peptidyl citrulline being reverted back to arginine (12). Therefore, citrullination is likely to accumulate in long lived proteins, such as histones, and prevent the target residues from undergoing electrophilic damage such as glycation (29). Based on this new deglycation activity of PAD4, it could potentially play a regulatory role in rescuing non-enzymatic damage and downstream changes in chromatin structure and function.

Another important implication of this finding is the three-way metabolic crosstalk between glycation, citrullination and methylation that compete for the same sites on histones (Figure 4). All these modifications contribute to chromatin architecture regulation and are directly correlated with the accessibility of the metabolites generated from the associated pathways: glycation with sugar glycolysis (30), citrullination with calcium homeostasis (12) and methylation with S-adenosyl methionine (SAM) metabolism (31), suggesting that this crosstalk is influenced by diet, metabolic state and the cellular microenvironment (32, 33). The balance between MGO, Ca^2+^ and SAM can be regulated through multiple processes including endoplasmic reticulum (ER) stress, reactive oxygen species (ROS) and mTOR signalling, providing an additional potential link to changes in gene expression (34-36). Together with our previous identification of the interrelationship between glycation and methylation (9, 10), this work suggests a three-way crosstalk (Figure 4) and new insights into the link between metabolism, epigenetics, and disease (37, 38).

**Figure 4.**
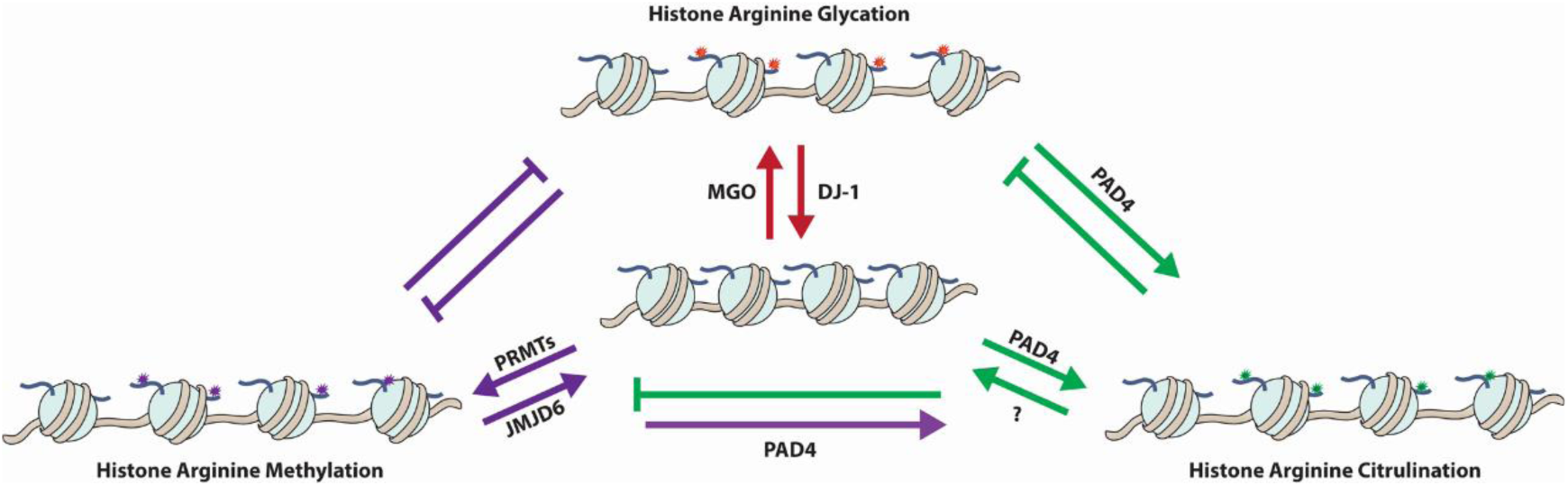
Three-way crosstalk of histone glycation, citrullination and methylation.

## Conclusions

Here we report an unprecedented dual function of PAD4 in antagonizing MGO-induced histone glycation. PAD4 induces changes in chromatin architecture by regulating not only charges of arginine residues but also the MGO-glycation level on histones. This unique function together with the overexpression of PAD4 in breast tumors provide insights into a potential mechanism for its function in cancer cells and understandings of the correlation between metabolism and cancer epigenetics.

## Materials and Methods

Materials, methods and supplementary results (protein expression and purification, *in vitro* and *in vivo* assays, *etc.*) are summarized in *SI Appendix, SI Materials and Methods, Supplementary Results*.

## ACKNOWLEDGMENTS

We would like to thank Prof. Paul Thompson at UMass Medical School for generously sharing the PAD plasmids and the detailed protocol for protein purification. We would also like to thank Prof. Minkui Luo at MSKCC for valuable discussions and critical reading of the manuscript. Work in the David lab is supported by R21 DA044767, CCSG core grant P30 CA008748, SPORE P50-CA192937 from the National Institutes of Health. In addition, work in the lab is supported by the Tri-institutional Therapeutic Discovery Institute, the Mr. William H. Goodwin and Mrs. Alice Goodwin and the Commonwealth Foundation for Cancer Research and the Center for Experimental Therapeutics at MSKCC, the Pershing Square Sohn Cancer Alliance and Cycle for Survival. A.O. is supported by the National Science Foundation Graduate Research Fellowship (Grant Number 2016217612) and the Chemical-Biology Interface training grant (NIH T32 GM115327-Tan). Y.D. is a Josie Robertson Young Investigator.

