## Supplemental Information for "Protein Arginine Deiminase 4 Antagonizes Methylglyoxal-induced Histone Glycation"

[3]Department of Pharmacology, Weill Cornell Medical College, New York, NY

### General materials and methods

UV spectrometry was performed on NanoDrop 2000c (Thermo Scientific). Biochemicals and media were purchased from Fisher Scientific or Sigma-Aldrich Corporation unless otherwise stated. T4 DNA ligase, DNA polymerase and restriction enzymes were obtained from New England BioLabs. PCR amplifications were performed on an Applied Biosystems Veriti Thermal Cycler using either Taq DNA polymerase (Vazyme Biotech) for routine genotype verification or Phanta Max Super-Fidelity DNA Polymerase (Vazyme Biotech) for high-fidelity amplification. Site-specific mutagenesis was performed according to the standard procedure of the QuickChange Site-Directed Mutagenesis Kit purchased from Stratagene (GE Healthcare) or Mut Express II (Vazyme Biotech). Primer synthesis and DNA sequencing were performed by Integrated DNA Technologies and Genewiz, respectively. PCR amplifications were performed on a Bio-Rad T100TM Thermal Cycler. Centrifugal filtration units were purchased from Millipore, and MINI dialysis units purchased from Pierce. Size exclusion chromatography was performed on an AKTA FPLC system from GE Healthcare equipped with a P-920 pump and UPC-900 monitor. Sephacryl S-200 columns were obtained from GE Healthcare. All the western blots were performed using the primary antibodies annotated in Supplementary Table 1 and fluorophore-labeled secondary antibodies annotated in Supplementary Table 2 following the protocol recommended by the manufacture. Blots were imaged on Odyssey CLx Imaging System (Li-Cor). Amino acid derivatives and coupling reagents were purchased from AGTC Bioproducts. Dimethylformamide (DMF), dichloromethane (DCM), triisopropylsilane (TIS) were purchased from Fisher Scientific and used without further purification. Hydroxybenzotriazole (HOBt) and O-(benzotriazol-1-yl)-N,N,N′,N′-tetramethyluronium hexafluorophosphate (HBTU) were purchased from Fisher Scientific. Trifluoroacetic acid (TFA) was purchased from Fisher Scientific. N,Ndiisopropylethylamine (DIPEA) was purchased from Fisher Scientific. Analytical reversed-phase HPLC (RP-HPLC) was performed on an Agilent 1200 series instrument with an Agilent C18 column (5 μm, 4 × 150 mm), employing 0.1% TFA in water (HPLC solvent A), and 90% acetonitrile, 0.1% TFA in water (HPLC solvent B) as the mobile phases. Analytical gradients were 0-70% HPLC buffer B over 45 min at a flow rate of 0.5 mL/min, unless stated otherwise. Preparative scale purifications were conducted on an Agilent LC system. An Agilent C18 preparative column (15-20 μm, 20 × 250 mm) or a semi-preparative column (12 μm, 10 mm × 250 mm) was employed at a flow rate of 20 mL/min or 4 mL/min, respectively. HPLC Electrospray ionization MS (HPLC-ESI-MS) analysis was performed on an Agilent 6120 Quadrupole LC/MS spectrometer (Agilent Technologies). All the immunoblotting experiments in this research were done at least in triplets. The error bars in all the figures represent the standard deviation from three different experiments.

### Recombinant histone expression and purification

Recombinant human histones H2A, H2B, H3.2, and H4 were expressed in *E. coli* BL21 (DE3) or *E. coli* C41 (DE3), extracted by guanidine hydrochloride and purified by flash reverse chromatography as previously described[1]. The purified histones were analyzed by RP-LC-ESI-MS: H2A $[M + H]^+$ observed, 13964.85 Da; expected, 13964.3 Da; H2B $[M + H]^+$ observed, 13758.79 Da; expected, 13758.9 Da; H3.2 $[M + H]^+$ observed, 15256.95 Da;

expected, 15256.8 Da; H4 $[M + H]^+$ observed, 11236.80 Da; expected, 11236.1 Da.

**Preparation of histone octamer and '601' DNA**

Octamers were prepared as previously described[1]. Briefly, recombinant histones were dissolved in unfolding buffer (20 mM Tris-HCl, 6M GdmCl, 0.5mM DTT, pH 7.5), and combined with the following stoichiometry: 1.1 eq. H2A, 1.1 eq. H2B, 1 eq. H3.2, 1 eq. H4. The combined histone solution was adjusted to 1 mg/mL concentration transferred to a dialysis cassette with a 7000 Da molecular cutoff. Octamers were assembled by dialysis at 4 °C against 3 × 1 L of octamer refolding buffer (10 mM Tris-HCl, 2 M NaCl, 0.5mM EDTA, 1 mM DTT, pH 7.5) and subsequently purified by size exclusion chromatography on a Superdex S200 10/300 column. Fractions containing octamers were combined, concentrated, diluted with glycerol to a final 50% v/v and stored at -20 °C. The 147-bp 601 DNA fragment was prepared by digestion from a plasmid containing 30 copies of the desired sequence (flanked by blunt EcoRV sites on either site), and purified by PEG-6000 precipitation as described before[2].

**Mononucleosome assembly**

The mononucleosome assembly was performed according to the previously described salt dilution method with slight modification[3]. Briefly, the purified wild-type octamers were mixed together with 601 DNA (1:1 ratio) in a 2 M salt solution (10 mM Tris pH 7.5, 2 M NaCl, 1 mM EDTA, 1 mM DTT). After incubation at 37 °C for 15 min, the mixture was gradually diluted (9 × 15 min) at 30 °C by dilution buffer (10 mM Tris pH 7.5, 10 mM NaCl, 1 mM EDTA, 1 mM DTT). The assembled mononucleosomes were concentrated and characterized by native gel electrophoresis (5% acrylamide gel, 0.5 × TBE, 120 V, 40 min) using ethidium bromide (EtBr) staining.

**Nucleosomal array assembly**

Dodecameric repeats of the 601 sequence separated by 30-bp linkers were produced from pWM530 using EcoRV digestion and PEG-6000 precipitation according to the published procedure[4]. Homotypic dodecameric arrays were assembled from purified octamers and recombinant DNA in the presence of buffer DNA (MMTV) by salt gradient dialysis as previously described[6]. The resulting arrays were purified and concentrated using $Mg^{2+}$ precipitation at 4 °C[3].

**Expression of recombinant PAD4**

The pGEX-PAD4 plasmid was a kind gift from Prof. Paul Thompson (UMass Medical School). The GST-tagged PAD4 protein was expressed in *E. coli* Rosetta (DE3) cells with an overnight IPTG induction at 16 °C. The bacterial pellet was lysed by sonication and lysate cleared by centrifugation at 12,000 r.p.m. for 30 min. Lysate was loaded on GSTrap HP Column (GE Healthcare) and eluted on AKTA FPLC (GE Healthcare) by gradient L-glutathione (reduced, Sigma). The GST tag was cleaved by Precission Protease overnight during dialysis, and the cleaved proteins was purified by reverse GSTrap HP Column and size exclusion chromatography on AKTA FPLC. Purified recombinant proteins were analyzed by SDS-PAGE, and concentrated using stirred ultrafiltration cells (Millipore) according to the manufacturer's protocol. The concentration of each protein was determined using 280 nm wavelength on a NanoDrop 2000c (Thermo Scientific).

**Peptide synthesis**

Standard Fmoc-based Solid Phase Peptide Synthesis (FmocSPPS) was used for the synthesis of peptides in this study. The peptides were synthesized on ChemMatrix resins with Rink Amide to generate C-terminal amides. Peptides were synthesized using manual addition of the reagents (using a stream of dry $N_2$ to agitate the reaction mixture). For amino acid coupling, 5 eq. Fmoc protected amino acid were pre-activated with 4.9 eq. HBTU, 5 eq. HOBt and 10 eq. DIPEA in DMF and then reacted with the N-terminally deprotected peptidyl resin. Fmoc deprotection was performed in an excess of 20% (v/v) piperidine in DMF, and the deprotected peptidyl resin was washed thoroughly with DMF to remove trace piperidine. Cleavage from the resin and side-chain deprotection were performed with 95 % TFA, 2.5% TIS and 2.5% $H_2O$ at room temperature for 1.5 h. The peptides were then precipitated with cold diethyl ether, isolated by centrifugation and dissolved in water with 0.1 % TFA followed by RP-HPLC and ESI-MS analyses. Preparative RP-HPLC was used to purify the peptides of interest.

***In vitro* biochemical assays**

The PAD4 citrullination assays were performed in the buffer (pH 7.5) containing 50 mM Tris-HCl, 5 mM $CaCl_2$ and 2 mM DTT (freshly added). For free histone H3 citrullination, 50 μM H3 were treated with 5 μM PAD4 at 37 °C for 2 h, and were analyzed by sodium dodecyl sulfate polyacrylamide gel electrophoresis (SDS-PAGE) followed by western blot analysis. For nucleosome core particle (NCP) citrullination, 1 μM NCPs were treated with 0.1 μM PAD4 at 37 °C for 2 h. The citrullinated NCPs were analyzed by SDS-PAGE or native page electrophoresis followed by western blot analysis. Nucleosomal array citrullination assays were similarly prepared using 1 μM dodecameric arrays and 0.1 μM PAD4 with slight modification, that is, the concentration of $CaCl_2$ was reduced to 100 μM to prevent the arrays from precipitation.

For the NCP citrullination-glycation assays, the wild type or citrullinated NCPs (1 μM) were treated with 5 mM MGO (Sigma) in 1 × PBS buffer (pH 7.4) at 37 °C for 6 h (short treatment) or overnight (long treatment)[5]. The buffer exchange between citrullination and glycation assays was performed using 0.5 mL Centrifugal Filter (3K, Millipore) with a 120-fold v/v for the removal of MGO or $Ca^{2+}$ from the old reaction buffer systems. The co-incubation of NCPs, MGO and PAD4 (or deglycase DJ-1) were also performed under the same conditions at 37 °C overnight. The modified NCPs were analyzed by SDS-PAGE (without boiling or lyophilizing the sample) or native gel followed by western blot analysis. For the SDS page analysis, H3 was used as loading control, while '601' DNA was used as loading control (by ethidium bromide staining) in the native gel analysis.

For peptide deglycation assays, 2 mM of the peptide substrate was incubated with 10 mM MGO in 1 × PBS buffer (pH 7.4) at 37 °C for 30 min and then enriched by magnetic streptavidin beads (Thermo Fisher Scientific, 65602). After being washed by 1 × PBS buffer, the glycated peptide was eluted with 100 mM glycine buffer (pH 2.5) and then diluted by using citrullination buffer (Tris-HCl), followed by the incubation with 100 μM PAD4 at 37°C for 2 h. The elution and reaction buffers used in peptide deglycation were made with $H_2^{18}O$ (Sigma-Aldrich, 329878). The reactions were analyzed by dot blot and LC-MS.

**$MgCl_2$ precipitation of nucleosomal arrays**

The $MgCl_2$ precipitation of nucleosomal arrays was performed according to the published procedure[6]. Briefly, increasing concentrations of $MgCl_2$ were added to the nucleosomal arrays and the reaction was incubated on ice for 10 min, followed by a 10-minute 17,000 rcf spin at 4 °C. The A260 of the supernatant was measured and used to evaluate the fraction of soluble arrays.

**Expression of PAD4 in 293T cells**

The PAD4 gene was amplified from pGEX-PAD4 by PCR. The primers used are 5'-ATATGCGGCCGCTACCCATACGATGTTCCAGATTACGCTATGGCCCAGGGGACATTG-3' (*NotI*) and 5'-ATATGGTACCTCAGGGCACCATGTTCCACC-3' (*KpnI*). The purified PCR product was inserted into pCMV plasmid by restriction endonuclease cloning. The catalytically dead mutant PAD4-C645S plasmid was constructed by site-directed mutagenesis using 5'-GGAGGTGCACAGCGGCACCAACG-3' and 5'-CGTTGGTGCCGCTGTGCACCTCC-3' as primers. The HA-tagged PAD4 was overexpressed in HEK 293T cells using Lipofectamine 2000 Transfection Reagent (Thermo Fisher Scientific) according to the manufacturer's protocol. HEK 293T cells (ATCC) were cultured at 37 °C with 5% $CO_2$ in DMEM medium supplemented with 10% fetal bovine serum (FBS) (Sigma-Aldrich), 2 mM L-glutamine and 500 units $mL^{-1}$ penicillin and streptomycin. Cells were stimulated with 2 μM calcium ionophore (Sigma-Aldrich, A23187) for 60 min at 37 °C before lysis, and then the PAD4 expression was detected by western blot analysis with anti-PAD4 and anti-HA antibodies.

**Salt extraction of histones from cells**

The extraction of histones from cells was performed according to the previously described high salt extraction method[7]. Briefly, the cell lysis solution was prepared using extraction buffer (10 mM HEPES pH 7.9, 10 mM KCl, 1.5 mM $MgCl_2$, 0.34 M sucrose, 10% glycerol, 0.2% NP40, protease and phosphatase inhibitors to 1 × from stock). After spinning down, the pellet was extracted using a no-salt buffer (3 mM EDTA, 0.2 mM EGTA). After discarding the supernatant, the final pellet was extracted by using high-salt buffer (50 mM Tris pH 8.0, 2.5 M NaCl, 0.05% NP40) in 4 °C cold room for 1 h. After spinning down, the supernatant containing extracted histones was collected for further analyses.

**Tumor samples**

All cell-culture reagents were obtained from Thermo Fisher Scientific unless otherwise indicated. The cell lines for tumor xenografts were maintained at 37 °C and 5% $CO_2$ in humidified atmosphere. MCF7 was obtained from DSMZ, while CAMA-1 and BT474 were obtained from ATCC. The cells were grown in DMEM/F12 supplemented with 10% FBS, 100 μg/mL penicillin, 100 mg/mL streptomycin, and 4 mmol/L glutamine. All the cell lines tested negative for Mycoplasma and authenticated by short-tandem repeat (STR) analysis. Six-to-8-week-old nu/nu athymic BALB/c female mice were obtained from Harlan

Laboratories, Inc., and maintained in pressurized ventilated caging. All studies were performed in compliance with institutional guidelines under an Institutional Animal Care and Use Committee-approved protocol (MSKCC#12–10–016). MCF7 and CAMA-1 xenograft tumors were established in nude mice by subcutaneously implanting 0.18-mg sustained release 17β-estradiol pellets with a 10 g trocar into one flank followed by injecting $1 \times 10^7$ cells suspended 1:1 (volume) with reconstituted basement membrane (Matrigel, Collaborative Research) on the opposite side 3 days afterward. The clinical samples (MSKCC set) used in this study were obtained from the Biobank of MSKCC. The Patients with breast cancer and either recurrence of disease after receiving adjuvant therapy or WHO-defined progression of metastatic disease on therapy were prospectively enrolled on an IRB approved tissue collection protocol (IRB#06-163). Informed consent was obtained from all patients. All patients underwent biopsy of at least a single site to document progressive disease. Mutational analysis of the metastatic biopsy was performed on fresh frozen specimens. Formalin fixed paraffin embedded (FFPE) blocks of the pretreatment primary tumor was obtained where possible for comparison. The presence of tumor, in both frozen samples and FFPE tissue sections, was confirmed by the study pathologist. Western blot analyses of PAD4 expression and histone citrullination were performed on fresh frozen specimens.

**Cell fractionation**

The cytosolic and nuclear fractions were prepared using NEPER Nuclear and Cytoplasmic Extraction Reagents (Thermo Scientific) according to the manufacturer's protocol. Histones were extracted from the pellet using high salt extraction protocol as described above[7]. Histone extraction from tumor xenografts and patient samples followed a similar protocol, with slight variation, where tumors homogenized by mild sonication prior to extraction. Purity of fractionation was evaluated using the following antibodies: anti-Actin (cytosol), anti-MEK ½ (nucleoplasm) and anti-H3 (chromatin).

**Pulse-chase experiments**

293T cells were treated with a gradient of MGO for 12 h before the medium was changed to MGO-free DMEM.[5] Cells were cultured for an additional 6 h, after which they were transfected with pCMV-PAD4 plasmid. After overnight incubation, the cells were harvested and cytosolic and histone fractions were prepared as described above. Samples were separated on a single SDS-PAGE, transferred to a PVDF membrane and blotted with the indicated antibodies.

**Micrococcal nuclease (MNase) digestion assay**

The MNase digestion assay was performed according to the previously described method with slight modification.[8] In brief, cell pellets were lysed in a hypotonic buffer (10 mM Tris-HCl pH 7.4, 10 mM KCl, 15 mM $MgCl_2$) on ice for 10 min. Nuclei were pelleted by centrifugation and resuspended in MNase digestion buffer (50 mM Tris-HCl pH 7.9, 5 mM $CaCl_2$) supplemented with RNase and incubated at 37 °C for 30 min. The DNA was then pelleted again by centrifugation and resuspended in MNase digestion buffer supplemented with 100 µg/ml BSA and 40 IU MNase and incubated at room temperature for varying periods of time (0, 5, 10 and 20 min). The MNase reaction was quenched with quenching buffer (0.4 M NaCl, 0.2% (w/v) SDS, 20 mM EDTA) followed by centrifugation. The DNA was extracted and purified by standard procedures, and then analyzed by Tris-Borate-EDTA (TBE) gel electrophoresis.

**Cell viability assay**

Untreated and 24-hours PAD4-transfected (WT or C645S) 293T cells were cultured in a 96-well plate and treated with the indicated MGO concentrations for 12 h. Following the incubation, cell viability was evaluated using the Cell Counting Kit-8 (CCK-8, Sigma) according to the manufacturer's protocol. The relative cell viabilities were given by detecting the absorbance at 460 nm at each well. Each experiment was performed in triplicate.

**Inhibitor treatment to breast cancer cell lines**

The PAD4 inhibitor GSK484 (Sigma, SML1658; 10 μM) was added to the MCF7 cells' media 6 h prior to adding the corresponding concentrations of MGO. Cells were incubated for additional 12 h after which they were harvested and histones were extracted and analyzed as described above. Samples were separated on a single SDS-PAGE, transferred to a PVDF membrane and blotted with the indicated antibodies.

**Supplementary figures**

**Figure S1.** SDS-PAGE analysis of PAD4 purification. The lanes from left to right are protein marker (M), concentrated PAD4 stock used for the *in vitro* assays (1), PAD4 before concentration (2), cleaved PAD4 before reverse GST column purification (3) and purified PAD4 before tag cleavage (4), respectively.

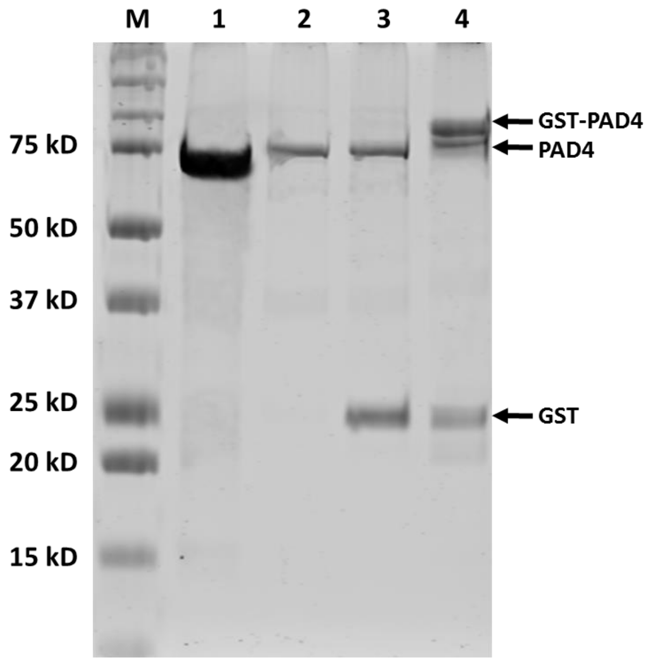

**Figure S2.** Differences of PAD4 enzymatic activity against distinct substrates (free histone H3, nucleosome core particles and nucleosomal arrays). Free histone H3, NCPs and nucleosomal arrays were treated with PAD4 at 37 °C for 2 h, and then separated by SDS-PAGE followed by western blot analysis.

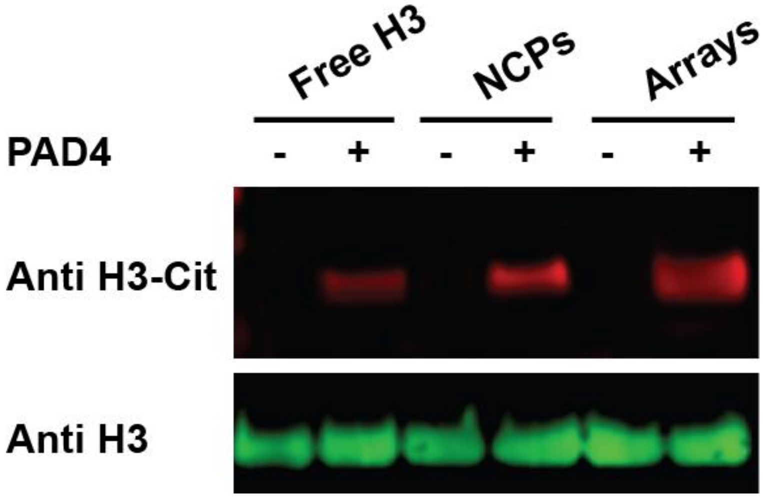

**Figure S3.** $Ca^{2+}$-dependent assays of PAD4 citrullination. The NCP citrullination reactions were performed in the presence or absence of $Ca^{2+}$ under 37 °C for 2 h, and then analyzed by native gel electrophoresis and western blot.

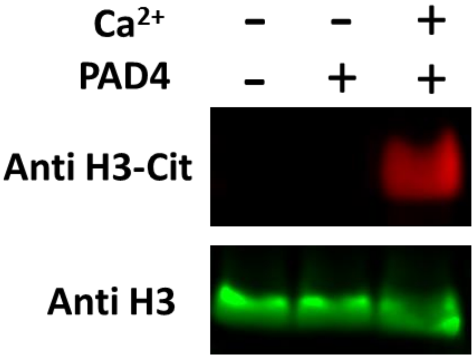

**Figure S4.** Dose-dependent NCP citrullination and glycation. NCPs were treated with gradient PAD4 under 37 °C for 2 h, followed by MGO treatment under 37 °C overnight, and then analyzed by native gel electrophoresis and western blot. The stoichiometry of enzyme and substrate is 0:20, 1:20, 2:20 and 4:20.

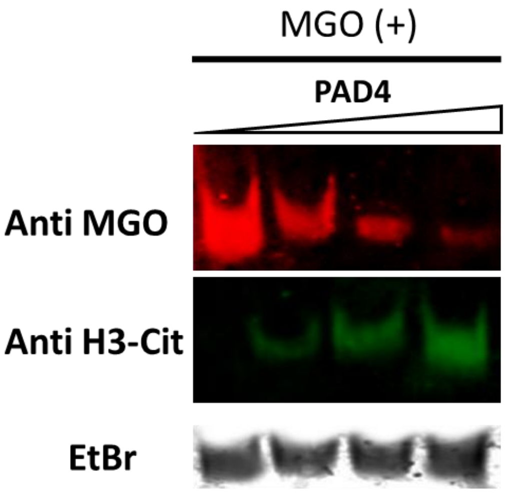

**Figure S5.** LC-MS analysis of MGO-induced glycation and PAD4-mediated deglycation on H3 N-terminal peptide substrate. The amino acid sequence of the peptide is ARTKQTARKSTGGKAPRK(Bio)A (calculated M=2237.3). (A) Ultraviolet absorption (λ=214 nm) of modified and unmodified peptides: **1** is the unmodified peptide (i), **2** is the citrullinated peptide by PAD4 (ii and iv), and **3** is the glycated peptide by MGO (iii). (B) Mass spectrometric analysis of corresponding peaks in A: **1** in i, M=2237.8; **2** in ii, M=2238.0; **3** in iii, M=2310.4; **2** in iv, M=2238.2.

**A**

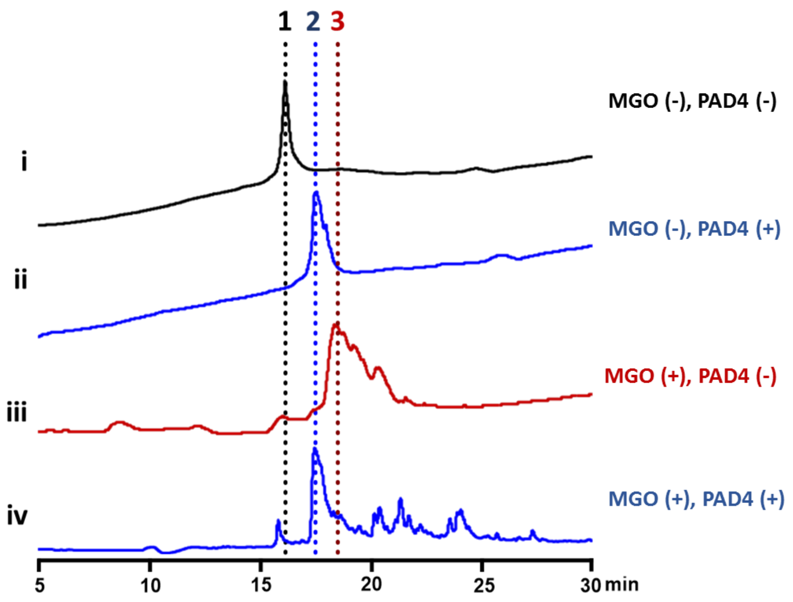


**B**

i

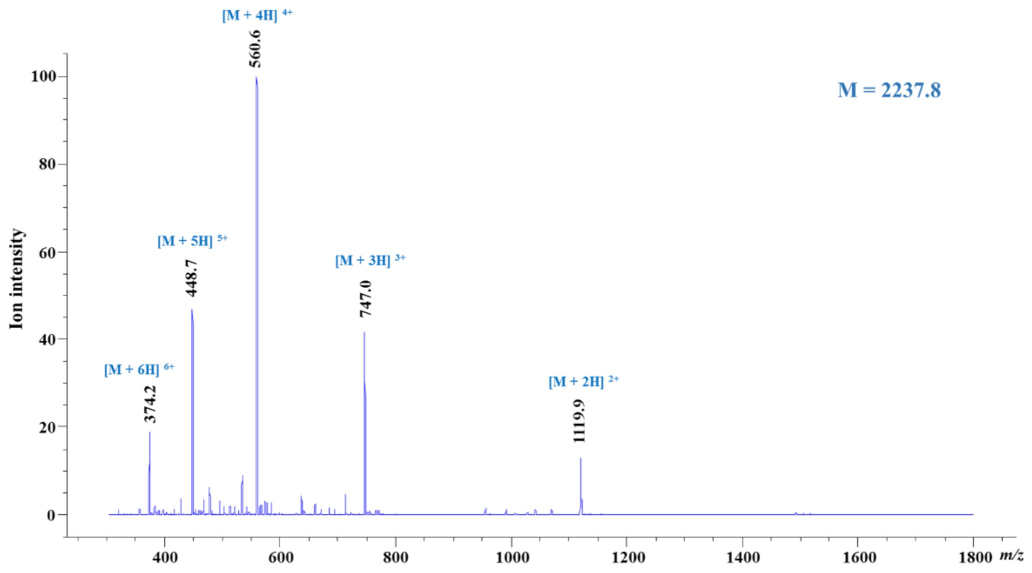

ii

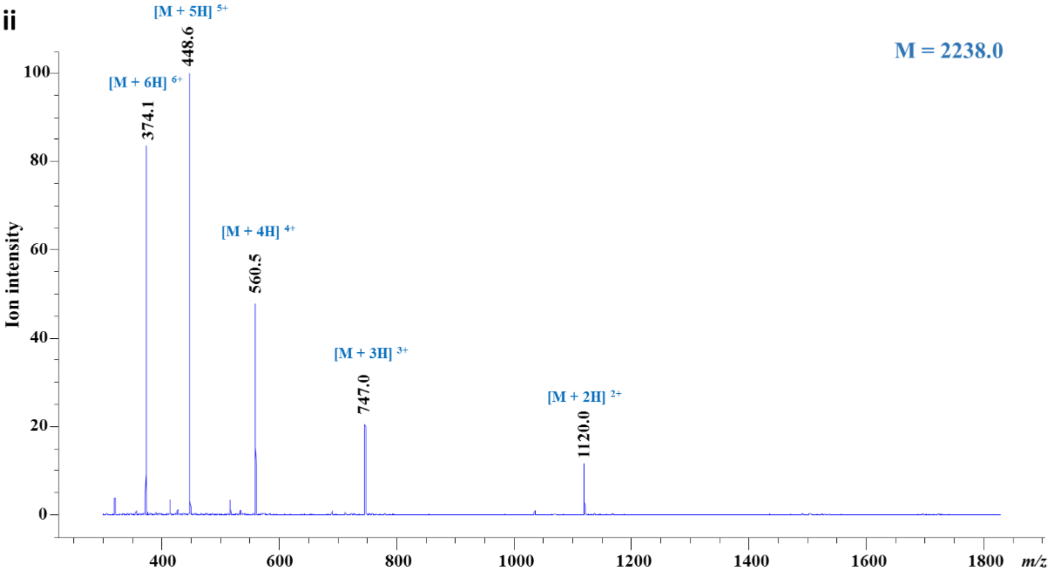
[M + 5H] 5+
448.6
[M + 6H] 6+
374.1
M = 2238.0
[M + 4H] 4+
560.5
[M + 3H] 3+
747.0
[M + 2H] 2+
1120.0
Ion intensity
m/z

iii

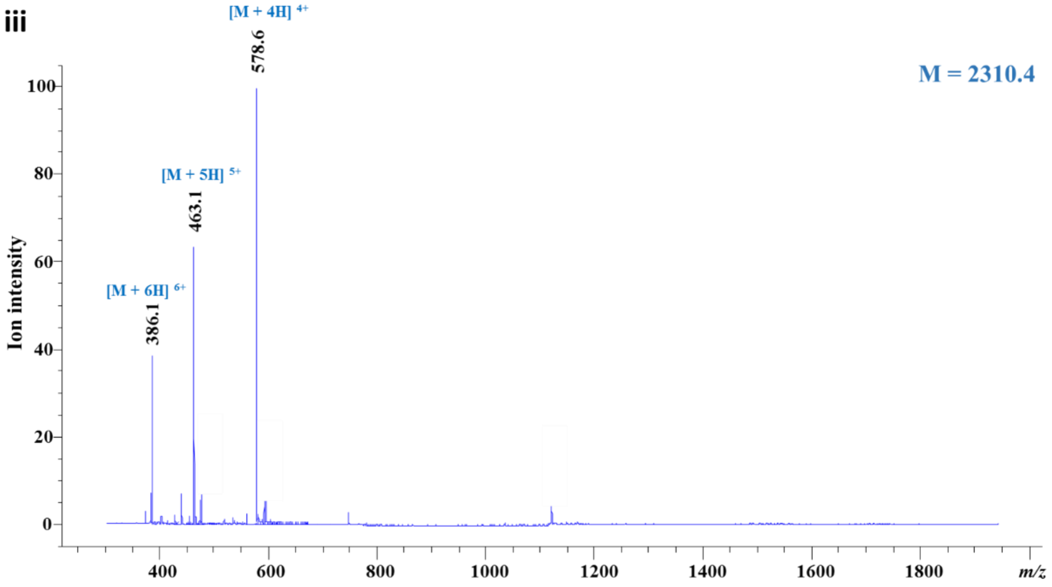
[M + 4H] 4+
578.6
M = 2310.4
[M + 5H] 5+
463.1
[M + 6H] 6+
386.1
Ion intensity
m/z

iv

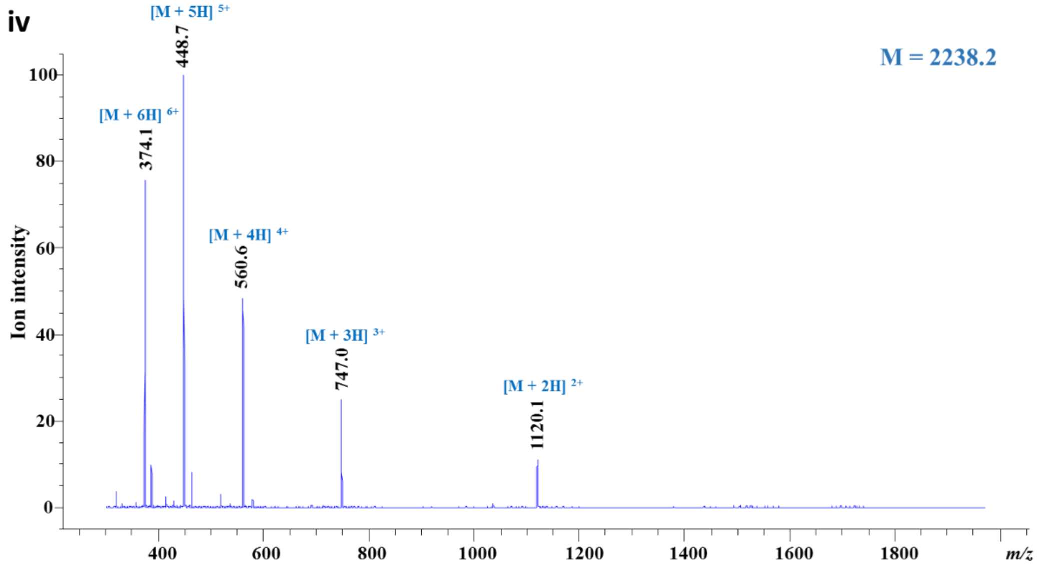
[M + 5H] 5+
448.7
[M + 6H] 6+
374.1
M = 2238.2
[M + 4H] 4+
560.6
[M + 3H] 3+
747.0
[M + 2H] 2+
1120.1
Ion intensity
m/z

**Figure S6.** LC-MS analysis of MGO-induced glycation and PAD4-mediated deglycation on H4 N-terminal peptide substrate. H4-R3(1-6) (calculated Mass=827.4) and H4-Cit3(1-6) (calculated Mass=828.4) were synthesized by SPPS as described in the Methods section and were used as substrate and product standard, respectively. MS analysis of the corresponding peaks in Fig. 2E: i) M=827.7, ii) M=828.5, iii) M=899.4, iv) M=830.4.

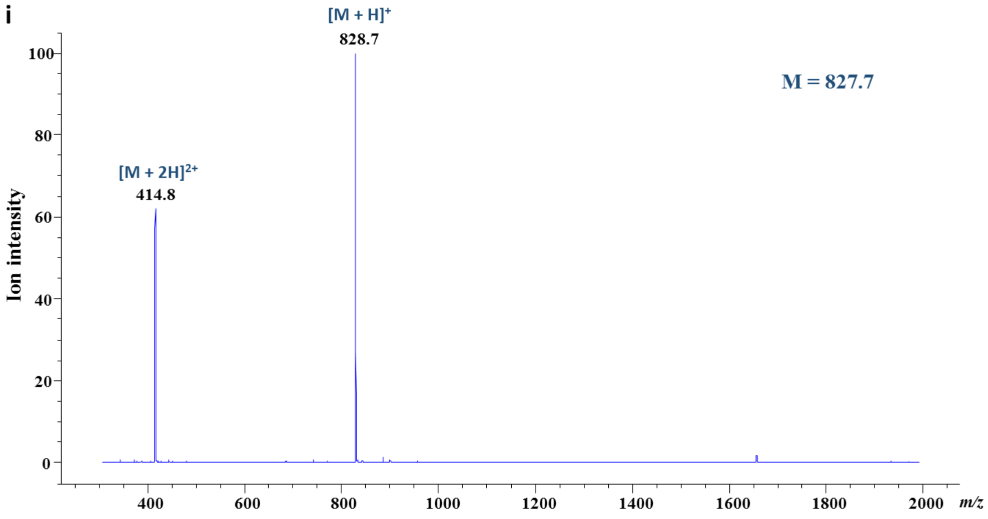


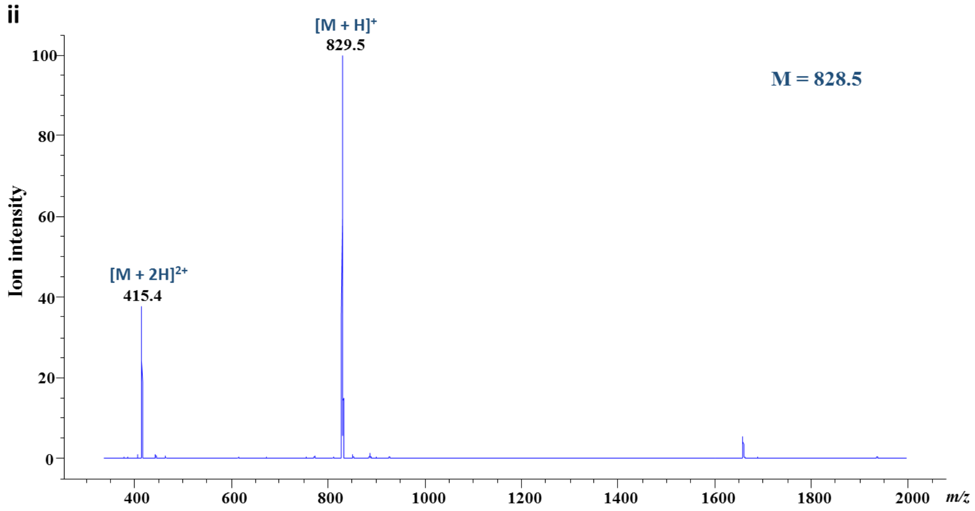

iii

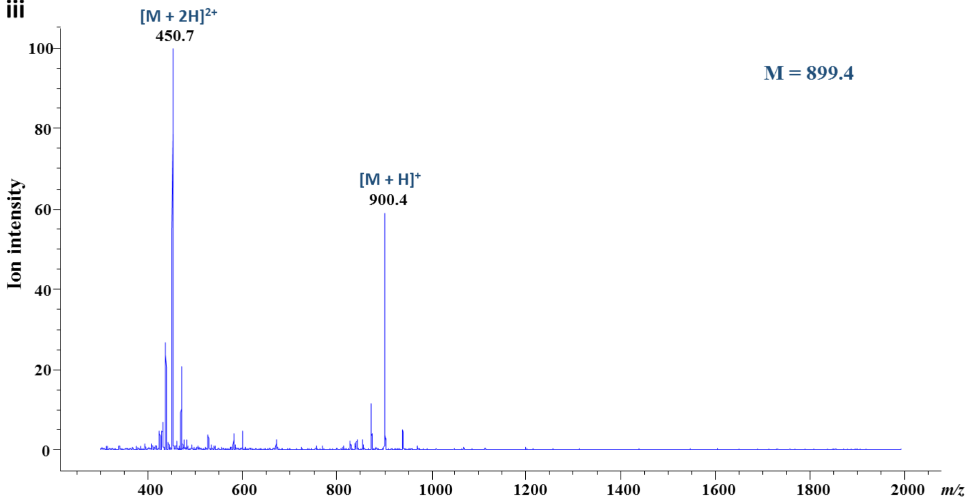
[M + 2H]2+
450.7
[M + H]+
900.4
M = 899.4
Ion intensity
100
80
60
40
20
0
400
600
800
1000
1200
1400
1600
1800
2000
m/z

iv

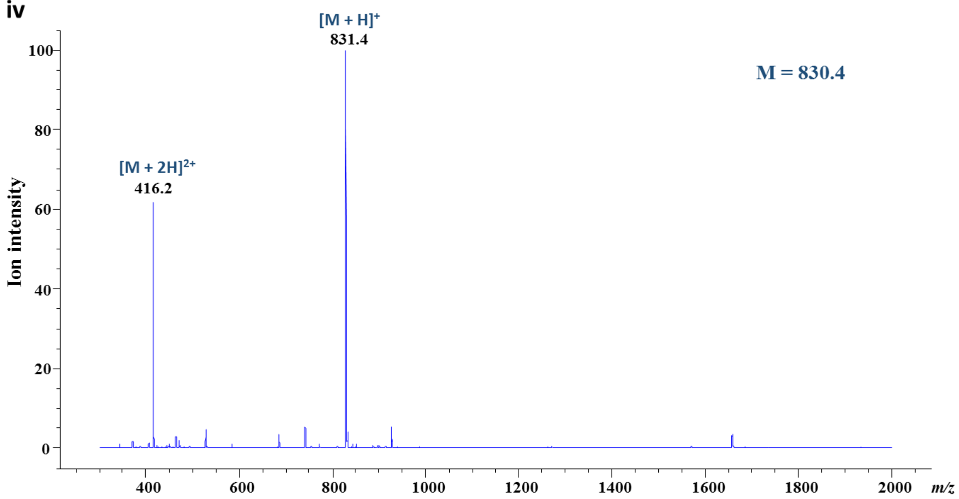
[M + H]+
831.4
[M + 2H]2+
416.2
M = 830.4
Ion intensity
100
80
60
40
20
0
400
600
800
1000
1200
1400
1600
1800
2000
m/z

**Figure S7.** Deglycase activities of DJ-1 and PAD4 on nucleosomal arrays. The arrays were pre-treated with 5 mM MGO for short (S, 6 h) and long period (L, 6 h + 12 h) in the presence or absence of enzymes. The enzymes were co-incubated with arrays and MGO (Co), incubated with glycated arrays after short treatment of MGO (S) or incubated with glycated arrays after long treatment of MGO (L).

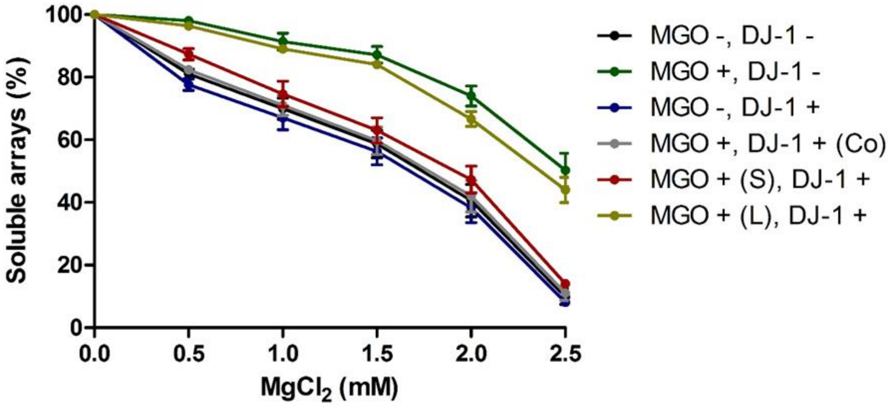

**Figure S8.** SDS-PAGE western blot analysis of extracted histones from wild type (left) and PAD4-overexpressing (right) 293T cells. Histones were extracted using salt extraction protocol described in supplementary method section. The gel was stained by Coomassie Brilliant Blue (CBB), and the histones were also transferred to a PVDF membrane and blotted with pan anti-citrulline antibody.

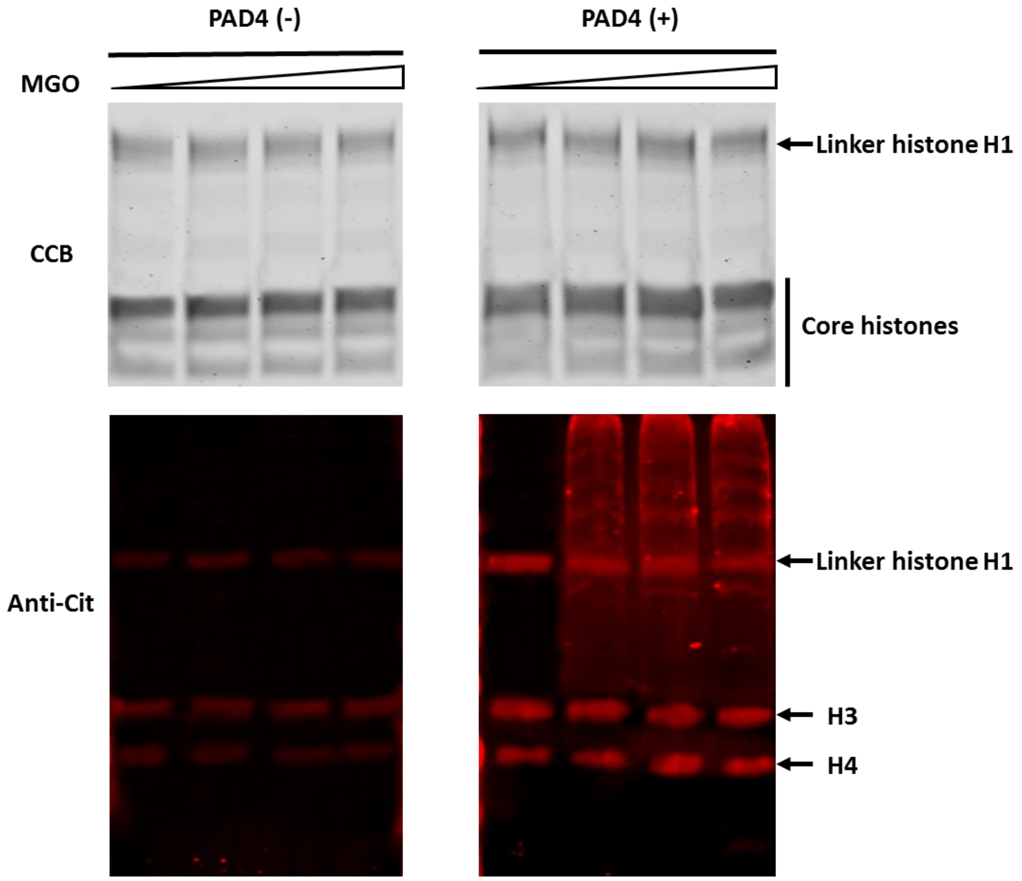

**Figure S9.** Pulse-chase analysis of histone glycation and PAD4-mediated deglycation. MGO pre-treated 293T cells were transfected with (right) or without (left) pCMV-PAD4 plasmid. The histones were extracted, separated by SDS-PAGE, transferred to a PVDF membrane and blotted with indicated antibodies.

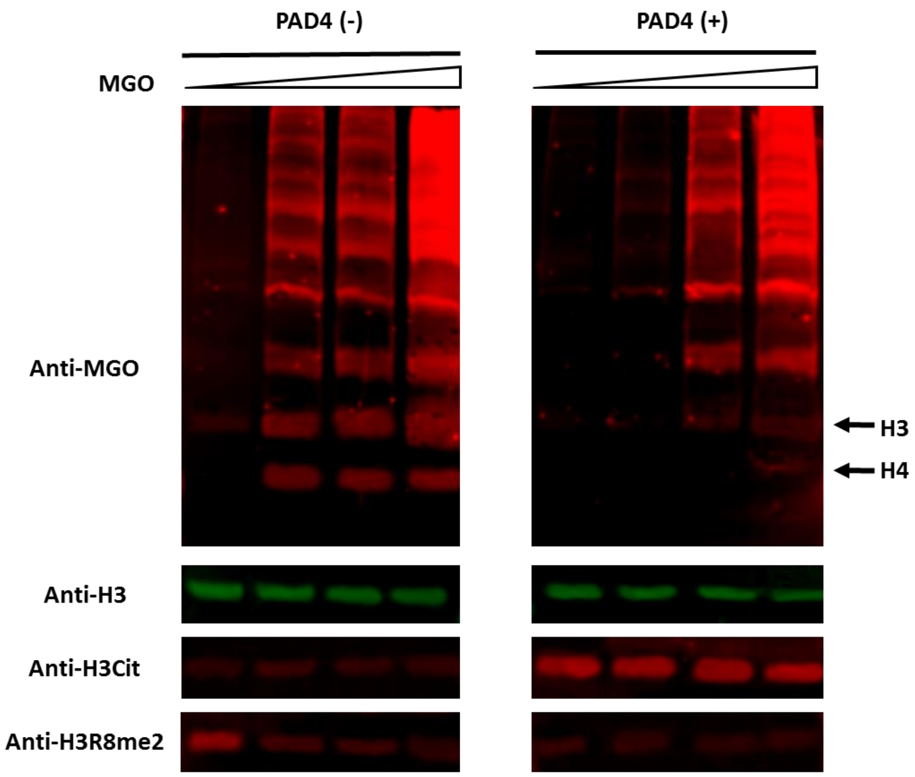

**Figure S10.** Sensitivity and compaction of MNase digested chromatin from WT and PAD4-overexpressed 293T cells with (left) and without 0.25 mM MGO (right) treatment. The chromatin were isolated from the cells and then digested by MNase at room temperature for varying periods of time (0, 5, 10 and 20 min). The DNA was extracted and purified by standard procedures, and then analyzed by Tris-Borate-EDTA (TBE) gel electrophoresis.

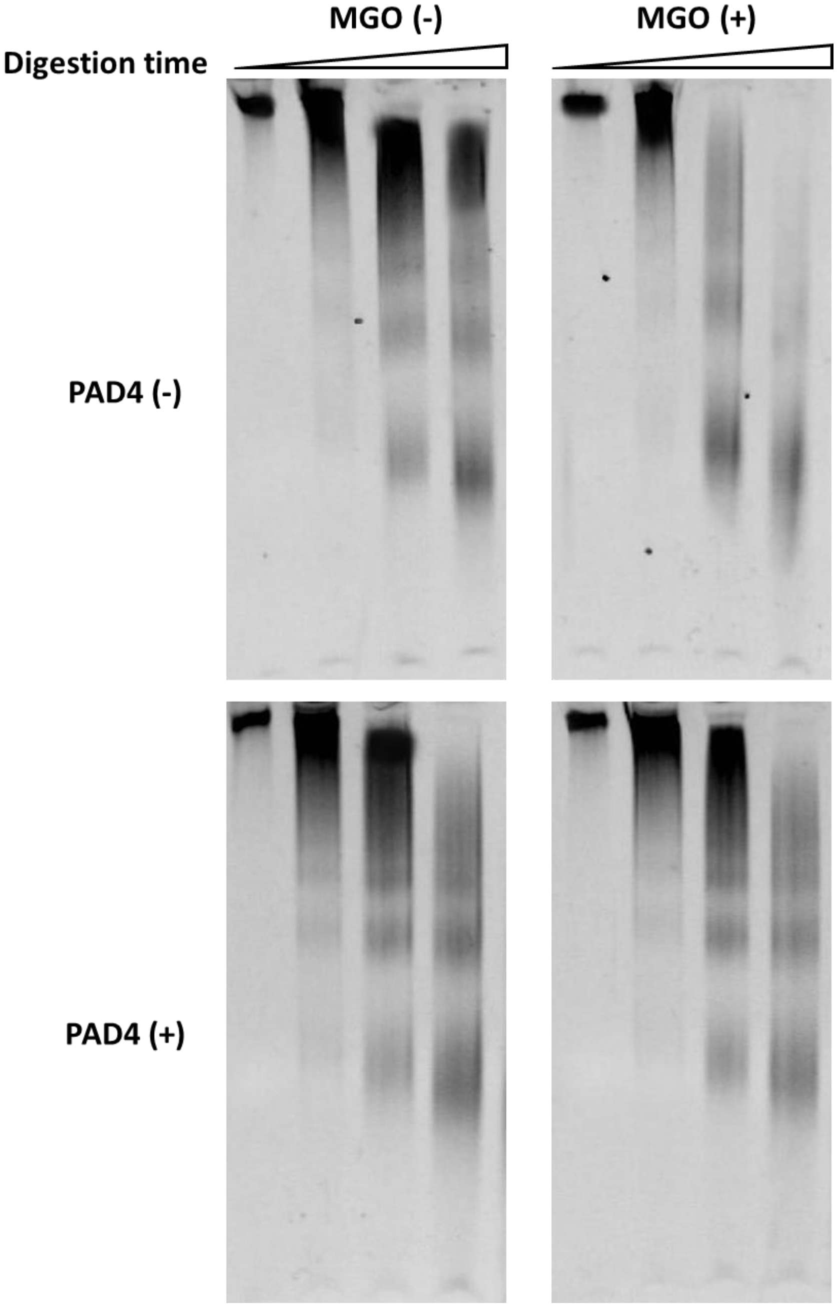

**Figure S11.** Cell viability of wild type (WT) and PAD4 (or PAD4-C645S)-overexpressed 293T cells non-treated (left) or treated with gradient MGO (right).

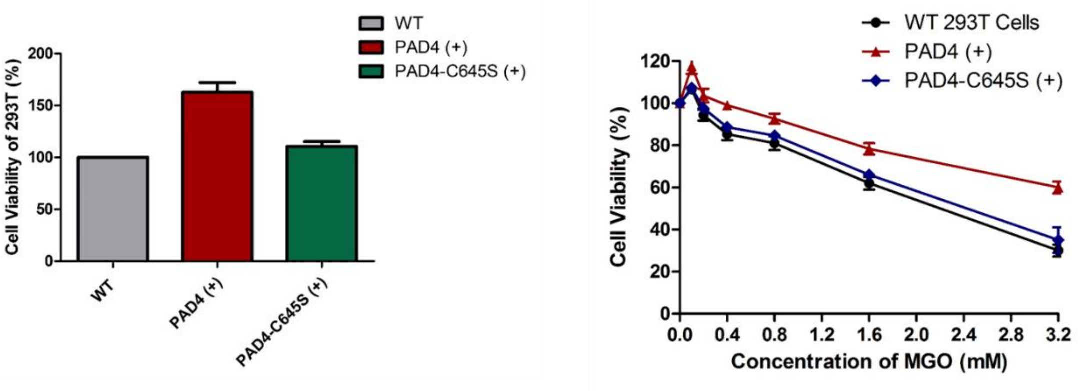

**Figure S12.** PAD4 inhibitor treatment to breast cancer cell line MCF7. MCF7 cells were pre-treated with PAD4 inhibitor GSK484 and then incubated with gradient MGO (0, 0.25, 0.5 and 1.0 mM). The histones were extracted, separated by SDS-PAGE, transferred to a PVDF membrane and blotted with indicated antibodies.

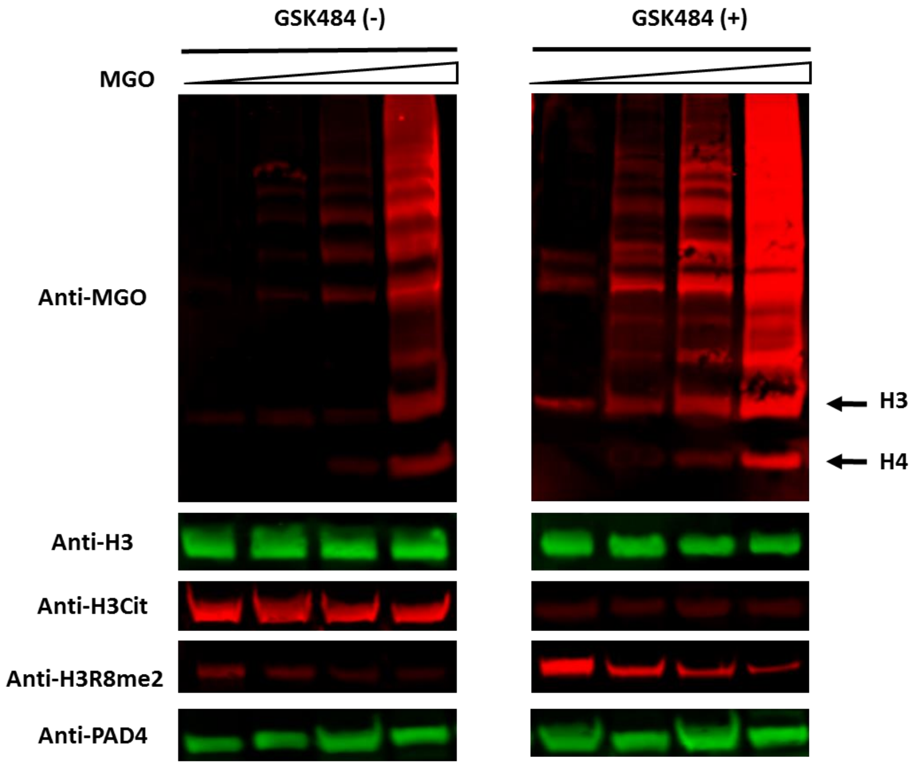

**Figure S13.** Quantification of selected immunoblotting (Figures 2B, 3A and S10) in this research. The error bars represent the standard deviation from three different experiments.

A

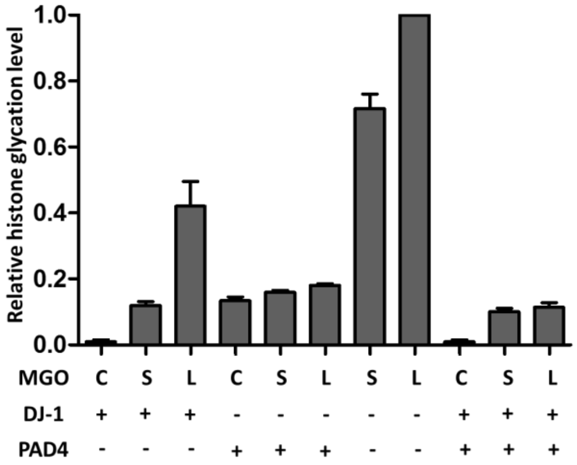


B

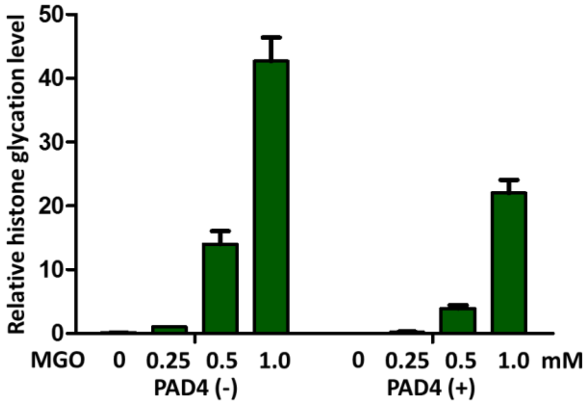


C

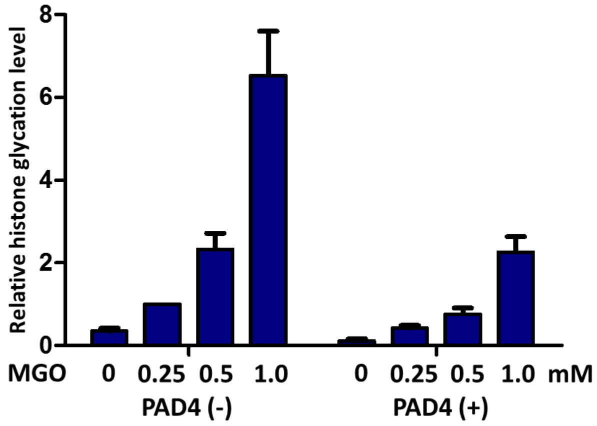

**Figure S14.** Proposed mechanisms and pathways of PAD4-mediated deglycation.

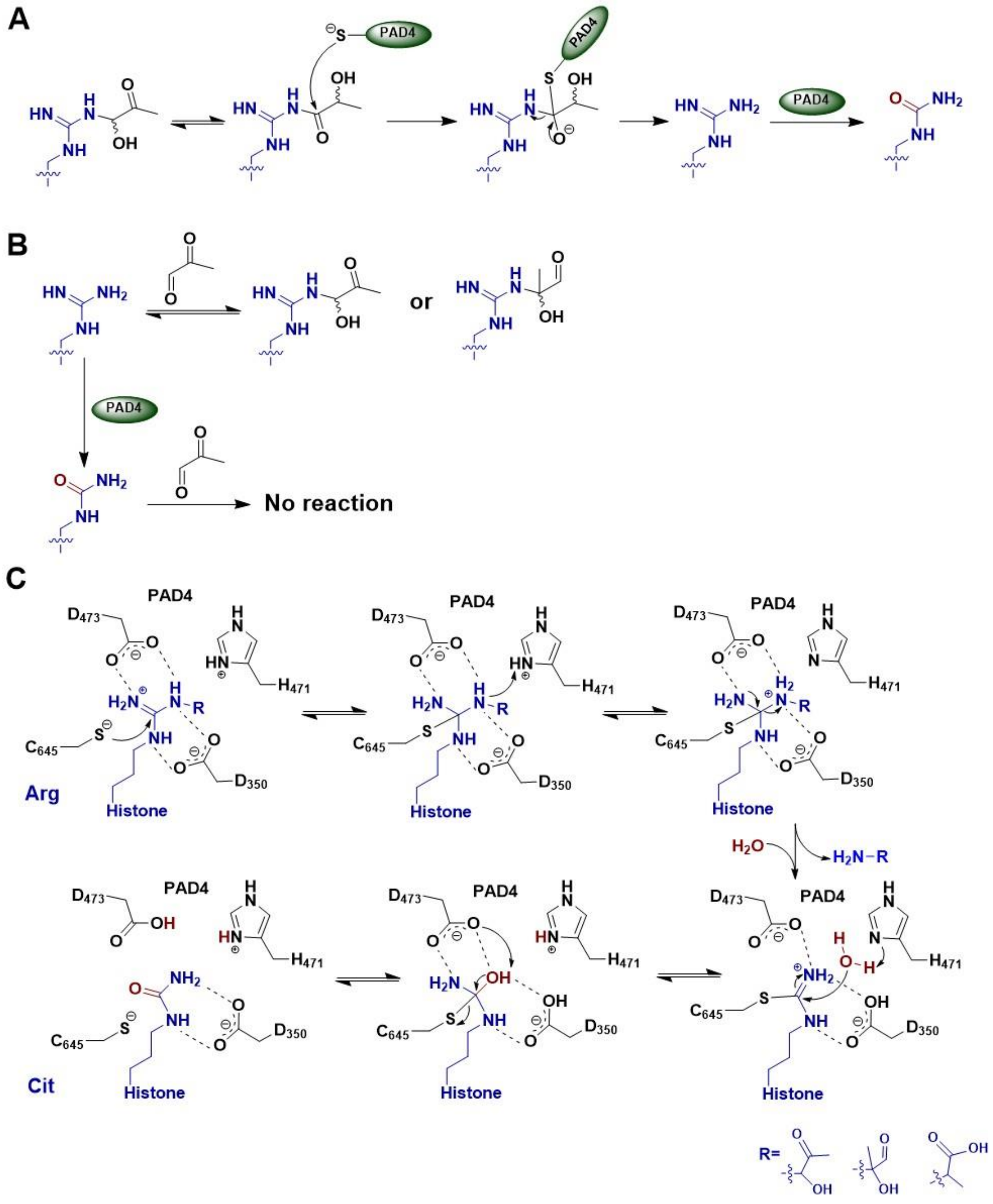

**Figure S15.** Schematic of MGO-induced glycation, the resulting species that can be recognized by the Anti-MGO antibody used in this study and the PAD4 substrates.

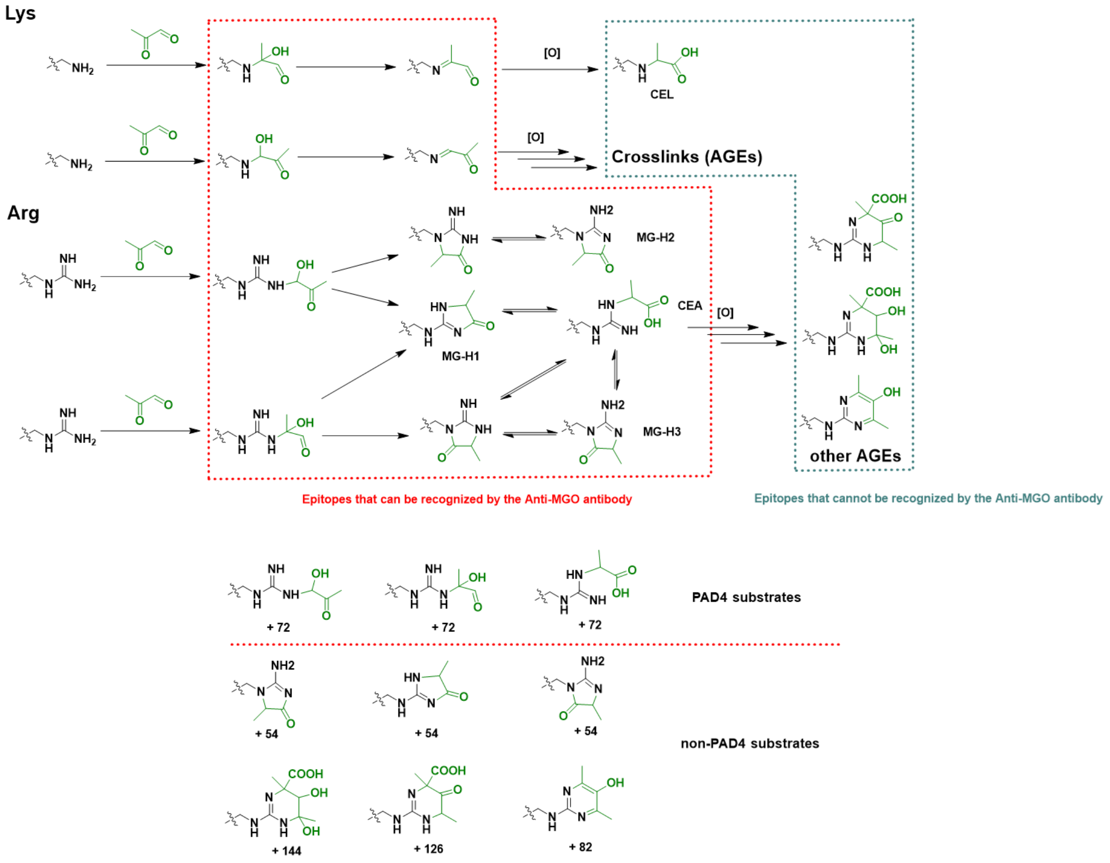

**Supplementary tables**

**Table S1.** Primary antibodies used in this manuscript.

| Host | Epitope | WB | Vendor |
|---|---|---|---|
| Mouse | Anti-MGO | 1: 500 | Cell Biolabs (STA-011) |
| Chicken | Anti-H3 | 1: 1000 | Abcam (ab134198) |
| Mouse | Anti-H3 | 1: 1000 | Abcam (ab10799) |
| Rabbit | Anti-H3Cit (R2, R8, R17) | 1: 1000 | Abcam (ab5103) |
| Rabbit | Anti-PAD4 | 1: 1000 | Abcam (ab50332) |
| Mouse | Anti-HA | 1:1000 | CST (3724S) |
| Mouse | Anti-Actin | 1: 1000 | CST (3700S) |
| Mouse | Anti-MEK 1/2 | 1: 1000 | CST (4694S) |
| Rabbit | Anti-H3R8Me2 | 1: 1000 | Abcam (ab194692) |
| Mouse | Anti-Citrulline | 1:1000 | Sigma (SAB5202274) |

**Table S2.** Secondary antibodies used in this manuscript.

| Host | Epitope | Label | Dilution | Vendor |
|---|---|---|---|---|
| Donkey | Anti-Chicken | IRDye 800CW | 1: 15000 | Li-Cor |
| Goat | Anti-Mouse | IRDye 680RD | 1: 15000 | Li-Cor |
| Goat | Anti-Mouse | IRDye 800CW | 1: 15000 | Li-Cor |
| Goat | Anti-Rabbit | IRDye 800CW | 1: 15000 | Li-Cor |
| Goat | Anti-Rabbit | IRDye 680RD | 1: 15000 | Li-Cor |
| - | Biotin | Atto 680-Streptavidin | 1:20000 | Sigma |
